# A *RUNX3* enhancer polymorphism associated with ankylosing spondylitis influences recruitment of Interferon Regulatory Factor 5 and factors of the Nucleosome Remodelling Deacetylase Complex in CD8+ T-cells

**DOI:** 10.1101/832840

**Authors:** Matteo Vecellio, Adrian Cortes, Sarah Bonham, Carlo Selmi, Julian C Knight, Roman Fischer, Matthew A Brown, B Paul Wordsworth, Carla J Cohen

**Affiliations:** Nuffield Department of Orthopaedics, Rheumatology and Musculoskeletal Sciences, University of Oxford, Oxford, UK; National Institute for Health Research Oxford Musculoskeletal Biomedical Research Unit, Oxford, UK; National Institute for Health Research Oxford Comprehensive Biomedical Research Centre, Botnar Research Centre, Nuffield Orthopaedic Centre, Oxford, UK; Nuffield Department of Clinical Neurosciences, Division of Clinical Neurology, John Radcliffe Hospital, University of Oxford, Oxford, UK; Wellcome Centre for Human Genetics, University of Oxford, Oxford, UK; Discovery Proteomics Facility, Target Discovery Institute, University of Oxford, Oxford, UK; Division of Rheumatology and Clinical Immunology, Humanitas Research Hospital, Rozzano, Milan, Italy; Guy’s, St Thomas’, King’s National Institute for Health Research Comprehensive Biomedical Research Centre

**Keywords:** spondyloarthropathy, genetic association, epigenetics, transcription factors

## Abstract

**Objectives:** To investigate the functional consequences of the single nucleotide polymorphism *rs4648889* in a putative enhancer upstream of the *RUNX3* promoter strongly associated with ankylosing spondylitis (AS).

**Methods:** The effects of *rs4648889* on transcription factor (TF) binding were tested by DNA pull-down and quantitative mass spectrometry. The results were validated by electrophoretic mobility gel shift assays (EMSA), Western blot (WB) analysis of the pulled-down eluates, and chromatin immuno-precipitation (ChIP)-qPCR.

**Results:** Several TFs showed differential allelic binding to a 50bp DNA probe spanning *rs4648889*. Binding was increased to the AS-risk A allele for IKZF3 (*aiolos*) in nuclear extracts from CD8+ T-cells (3.7-fold, p<0.03) and several components of the NUcleosome Remodeling Deacetylase (NuRD) complex, including Chromodomain-Helicase-DNA-binding protein 4 (3.6-fold, p<0.05) and Retinoblastoma-Binding Protein 4 (4.1-fold, p<0.02). In contrast, binding of interferon regulatory factor (IRF) 5 was increased to the AS-protective G allele. These results were confirmed by EMSA, WB and ChIP-qPCR.

**Conclusions:** The association of AS with *rs4648889* most likely results from its influence on the binding of this enhancer-like region to TFs, including IRF5, IKZF3 and members of the NuRD complex. Further investigation of these factors and RUNX3-related pathways may reveal important new therapeutic possibilities in AS.

## INTRODUCTION

Characterised by prominent axial skeletal involvement and enthesitis ankylosing spondylitis (AS) is the archetypal inflammatory spondyloarthropathy (SpA). It has a complex aetiology with more than a hundred genetic influences.[1, 2] Only a few of the associated single nucleotide polymorphisms (SNP) cause obvious functional amino acid substitutions, such as *rs11209026* that impairs signalling through the interleukin (IL) 23 receptor.[3] In contrast, most disease-associated SNPs probably exert a regulatory role on gene expression through epigenetic mechanisms, such as differential binding of transcription factors (TF) or microRNAs.[4] Here we report the extended effects on TF binding of a SNP (*rs4648889*) in a putative regulatory element (enhancer) upstream of the *RUNX3* gene, which is strongly associated with AS. RUNX3 is a key regulator of several lineage-specific developmental pathways, including T-cells, and has potential roles in infection, immunity and cancer.[5, 6] We previously showed that *rs4648889* influences *RUNX3* expression in CD8+ T-cells by altered binding of the interferon regulatory factor (IRF) 4 TF.[7] The effects of TF complexes bound to DNA may depend on the overall complement of components recruited to them. We therefore used a DNA pulldown approach combined with quantitative mass spectrometry to define the full range of interacting TF partners binding at *rs4648889* and to refine our previous observations relating to IRF4.[7, 8] Here we show that IKZF3 (the zinc finger protein *aiolos*) and several factors of the NUcleosome Remodeling Deacetylase (NuRD) complex (a gene repressor complex involved in chromatin remodelling) show differential allelic binding at *rs4648889* and that IRF5 is also involved.

## RESULTS

### SNP-based TF capture and analysis by label-free mass spectroscopy

Principal component analysis (PCA) showed well-separated clusters corresponding to the different *rs4648889* genotypes (Fig 1A). Significant differential binding was apparent for 38 proteins (false discovery rate <0.05) between the protective “G” allele and the AS-risk associated “A” allele (Fig 1B and Suppl Table 1). Reactome pathway analysis revealed significant enrichment for proteins involved in immunity, chromatin remodelling/histone deacetylation, RNA polymerase II transcription initiation and *RUNX3* regulation (Fig 1C). Unsupervised hierarchical clustering demonstrated distinct DNA/protein “interactome” profiles for these two alleles (Fig 1D).

**Figure 1.**
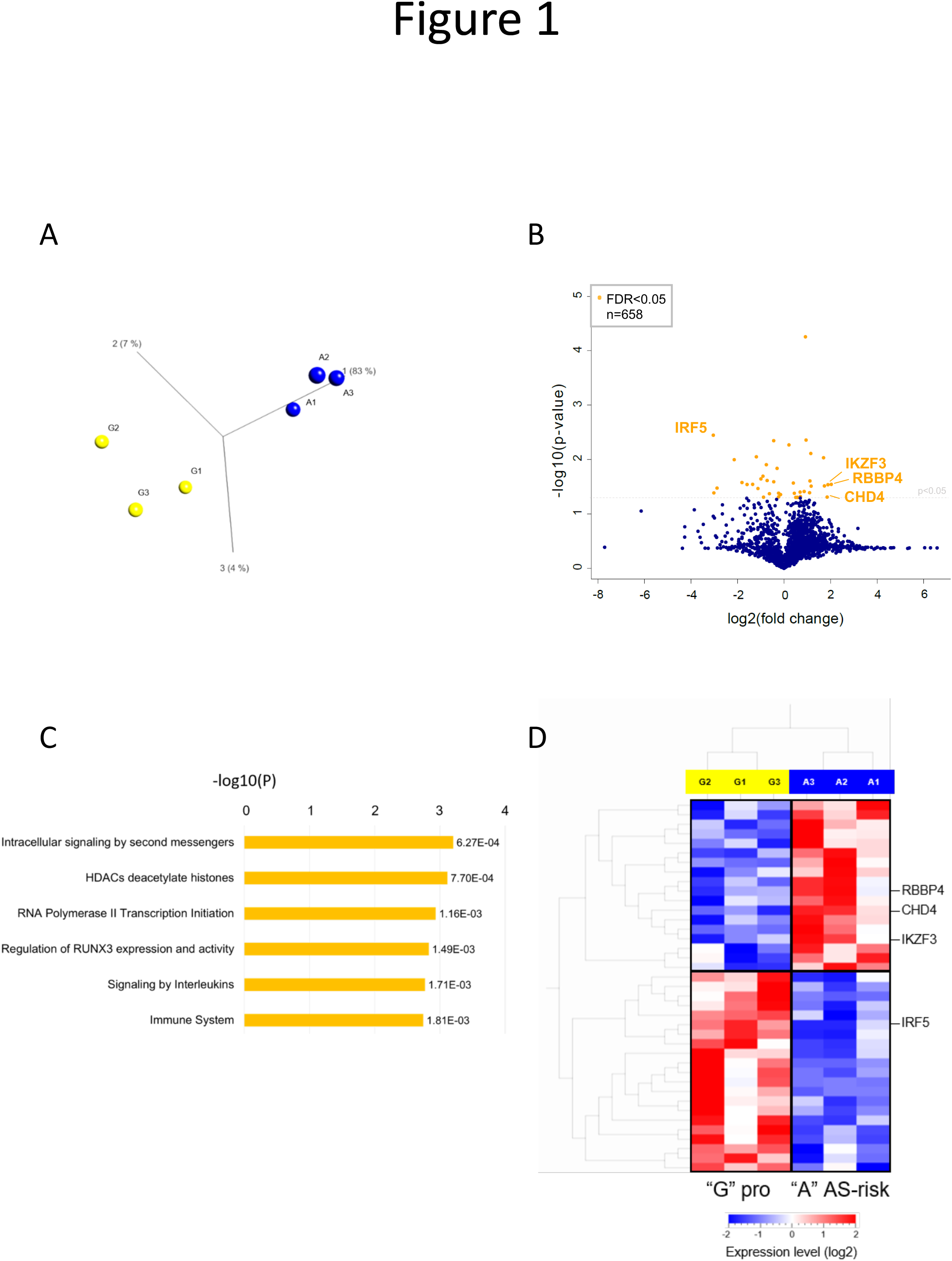
Identification of the proteins bound to the *rs4648889* locus with DNA capture assay/quantitative MS. (A) Principal component analysis (PCA) plot of the three replicates for the “G” protective allele vs the “A” AS-risk allele. (B) Volcano plot showing the complete set of 658 proteins identified in three different experiments. The orange dots represent the statistically significant proteins resulting from log2(fold change)/-log10(p-value). (C) Reactome pathway analysis of the statistically significant proteins identified. (D) Unsupervised hierarchical clustering of the statistically significant proteins (p<0.05) differentially quantified between the two alleles.

### Identification of IKZF3 and NuRD co-factors by Mass Spectrometry (MS)

Specific DNA-binding motifs encompassing *rs4648889* were found for IKZF3 and Chromodomain-Helicase-DNA-binding protein 4 (CHD4) (Fig 2A). Analysis of publicly available ChIP-seq data (ENCODE https://genome.ucsc.edu/ENCODE/) confirmed strong peak signals both for IKZF factors and CHD4 in the vicinity of *rs4648889*, in lymphoblastoid cell lines (Fig 2B). The pull-down experiments (Fig 2C and Suppl Table 2) identified all the previously described members of the NuRD repressor complex.[9] The label-free quantitation showed that IKZF3 was significantly more abundant with the AS-risk A allele compared to the protective G allele (3.7-fold, p<0.05). Binding to the A allele was also significantly increased for several components of the NuRD complex, including CHD4 (3.6-fold, p<0.05), Retinoblastoma-Binding Protein 4 (RBBP4, 4.1-fold, p<0.03), and Methyl-CpG-Binding Domain protein 2 (MBD2, 1.5-fold, p=0.05). This trend continued for other NuRD proteins, including Lysine–Specific histone Demethylase 1 (KDM1A) and the Histone Deacetylases HDAC1 and HDAC2 (non-significant increased binding to the A allele 2.4-, 16.1- and 16.2-fold, respectively).

**Figure 2.**
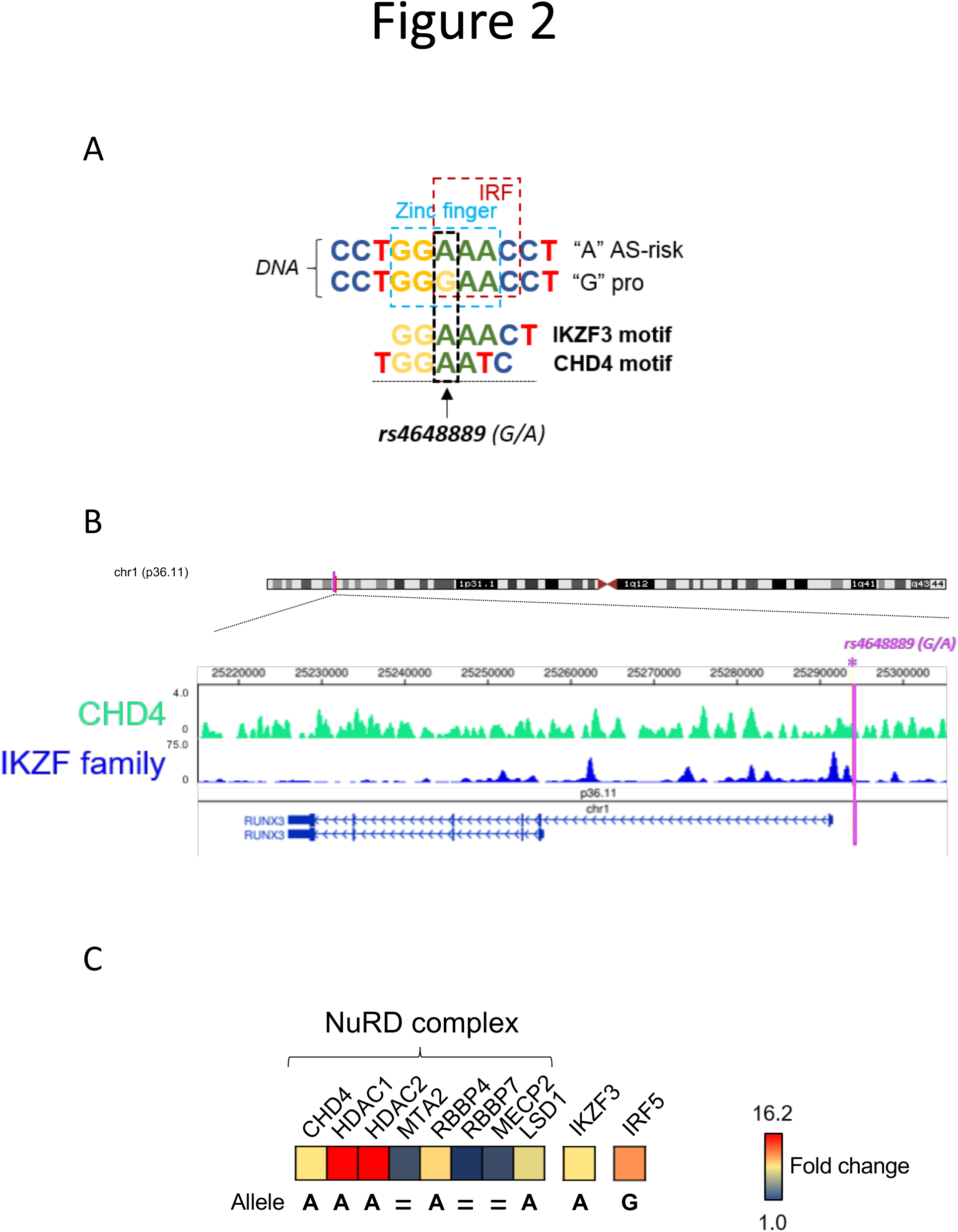
Identification of IKZF3, CHD4 and NuRD co-factors in binding the *rs4648889* encompassing genomic locus. (A) DNA binding motifs (from ENCODE – Factorbook [18]) of IKZF3 and CHD4 which overlap the *rs4648889* encompassing locus. (B) Epigenome browser database interrogation showing ChIP-seq peak signal for CHD4 and IKZF factors on GM12878 cell lines. Purple line locates *rs4648889* genetic variant. (C) Label-free quantitation of IKZF3 and NuRD complex factors (and IRF5) identified by MS. The graph shows differential binding expressed as fold change (blu= no difference, red=high difference).

### IKZF3-NuRD-RUNX3 pathway analysis

There was significant enrichment for protein-protein interactions among the differentially abundant NuRD-related proteins, IKZF3 and RUNX3 (p<7.4×10^−9^; Suppl Fig 2A), relating to transcription, DNA-binding and histone deacetylation (respectively p=6.3×10^−5^, p=6.1×10^−6^, p=3.6×10^−9^; Suppl Fig 1B) in the GO molecular functions and biological processes function (Suppl Fig 2B).

### Validation of differential binding for IKZF3, CHD4 and RBBP4

The MS results were validated by three further experimental approaches. First, Western blots (WB) of the pulled-down eluates confirmed the preferential binding of IKZF3, CHD4 and RBBP4 to the AS-risk A allele (Fig. 3A). Second, the impact of *rs4648889* on TF binding was confirmed in EMSA performed with nuclear extract (NE) from Jurkat T-cell line. The intensity of the DNA/NE complex was less with the AS-risk allele than the protective allele (G=68.1 ± 1.5 vs A=21.3 ± 3.7; p=0.002; n=3). A supershifted band more evident with the AS-risk A allele was induced by antibodies against IKZF3, CHD4 (Fig 3B, lanes 9 and 10) and RBBP4 (Fig 3D, lane 8) confirming the interaction of these TF with the 50bp sequence encompassing *rs4648889*. Third, allele-specific ChIP-qPCR of freshly isolated CD8+ T-cells from blood cones, heterozygous for *rs4648889* (3 independent experiments) showed enhanced relative enrichment for IKZF3, CHD4, RBBP4 (p=0.04, p=0.07 and p=0.06, respectively) to the AS-risk A allele (Fig 3E).

**Figure 3.**
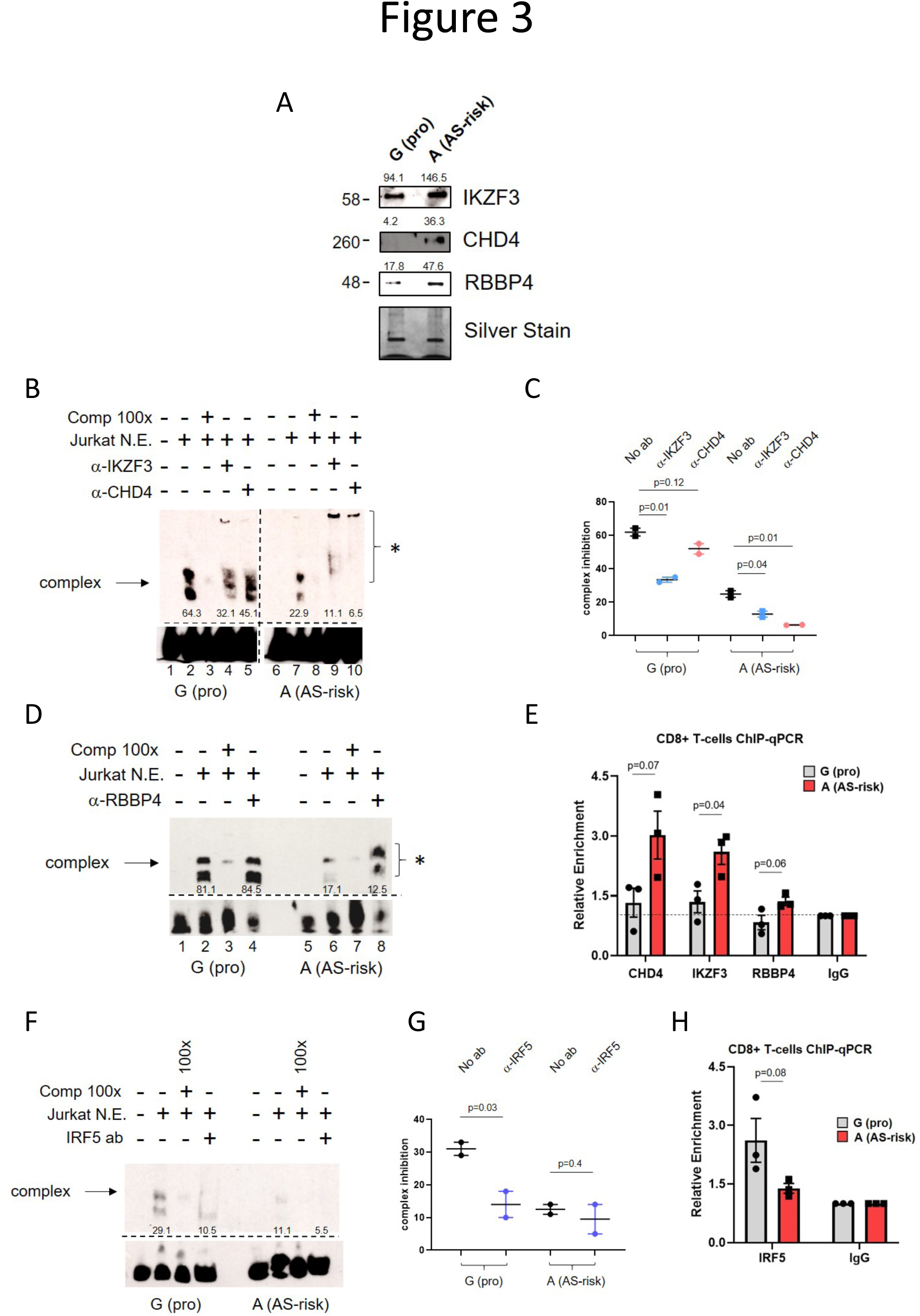
Validation of the factors identified. (A) Representative Western Blot on the pulled-down eluates used in MS experiments assessing the differential expression for IKZF3, CHD4 and RBBP4. The numbers above the bands show their quantification performed with Image J. (B) Representative EMSA (n=2) showing differential nuclear extract binding (complex indicated by arrow) after addition of Jurkat cell lysate (lanes 2 and 7). The 100-fold excess of unlabelled probes has been used as competitor (lanes 3 and 8). The involvement of IKZF3 and CHD4 was assessed by adding the corresponding antibody (lanes 4, 5, 9 and 10). Asterisk (*) indicates the presence of a super-shifted complex. The numbers below the bands represent the pixel intensity measured with Image J. (C) The graph shows the quantification of the inhibition of complex formation after addition of the antibodies (n=2) performed with Image J. (D) Representative EMSA (n=2) showing differential nuclear extract binding of RBBP4, after addition of the RBBP4 antibody (lanes 4 and 8). Asterisk (*) indicates super-shift. (E) The relative enrichment of CHD4, IKZF3 and RBBP4 was assessed with chromatin immunoprecipitation experiments (n=3) on CD8+ T cells (from buffy coat blood cones) heterozygous for *rs4648889*. P-value is indicated on top of the bars for each gene. Data are normalized on input and IgG is set as 1. (F) Representative EMSA (n=2) showing differential binding for IRF5, after addition of the antibody (lanes 4 and 8). In this case inhibition of the complex has been assessed. (G) The graph is showing the quantification of the complex inhibition after addition of IRF5 antibody (n=3) performed with Image J. (H) The relative enrichment of IRF5 was assessed with allelic-specific ChIP-qPCR on CD8+ T cells (from buffy coat blood cones) heterozygous for *rs4648889*.

### Interferon regulatory factor 5 (IRF5) preferentially binds to the G allele at *rs4648889*

IRF4 was not among the 38 proteins displaying significant differential allelic binding using this approach but IRF5 bound preferentially to the AS-protective G allele (fold change 8.2, p=0.003). IRF5 binding to *rs4648889* was therefore also further evaluated. EMSA revealed markedly greater binding of Jurkat NE to the G allele, which was significantly reduced by IRF5 antibody (Fig 3 F & G). ChIP-qPCR was also suggested allelic imbalance with a non-significant enhanced enrichment for the protective G allele (Fig 3H, p=0.08).

## DISCUSSION

We identified a complex network of TFs and chromatin regulatory proteins [10, 11] interacting with a putative cis-regulatory (enhancer) element upstream of the distal *RUNX3* promoter, which is implicated in the aetiology of AS. Many of these proteins exhibit differential allelic binding *in vitro* to a short DNA sequence in this region around *rs4648889*. In particular, IKZF3 (*aiolos*), a global regulator of chromatin architecture,[12] binds preferentially to the AS-risk allele. The Ikaros family of TFs (including *aiolos*) play important roles in lymphocyte development and are often associated with the NuRD complex, an ATP-dependent chromatin-remodelling complex involved in transcriptional repression.[13, 14] Our DNA pull-down approach identified most of the components of the NuRD complex active at this locus (CHD4, MBD2, RBBP4, HDAC1, HDAC2 and KDM1A), all of which exhibit the same preferential binding to the A allele. Results consistent with these were also obtained by WB, EMSA and ChIP-qPCR, suggesting that transcriptional repression [7] associated with the NuRD complex is implicated in the pathogenesis of AS perhaps by regulating *RUNX3* expression in CD8+ T-cells. Interestingly certain T-cell memory phenotypes are reliant on the recruitment of Eomesodermin (another TF encoded at the AS-associated *EOMES* locus)[1] that binds mostly *to RUNX3* bound enhancers, such as HDACs and CHD4.[15] Rheumatologists also have an interest in CHD4 because it is one of the target antigens for autoantibodies (anti-Mi2) in about 20 per cent of cases of dermatomyositis.[16] However, the relevance of these antibodies to the inflammatory processes of myositis or the malignancies commonly associated with dermatomyositis is unclear.

IRF5 was also among the 38 factors with significant differential allelic binding (p=0,003) but in this case the binding was significantly higher to the protective G allele. Previously we identified IRF4, another IRF family member, binding to this region [7] but we could not identify IRF4 in our pull-down experiments here. Overall our results suggest that both factors bind to the DNA region upstream *RUNX3* but suggest either differential activity between IRF4 and IRF5 or lower expression of IRF4, which was not therefore detectable by mass spectrometry. The consensus IRF motifs (GGAAC/GAAAC) overlap those for *CHD4* and *IKZF3* (*aiolos*) at *rs4648889* (Figure 2A), which raises the possibility of mutually antagonistic effects on DNA binding. IRF5 is of considerable interest in inflammatory states: it plays a central role in the induction of inflammatory cytokines, such IL6, IL12, IL23 and tumour necrosis factor α.[17] It is also a key factor in determining inflammatory macrophage phenotypes in response to stimulation by interferon γ and granulocyte-macrophage stimulating factor, which is now thought to be a key factor in the development of AS.[18] IRF5 is also strongly associated genetically with other inflammatory disorders, including rheumatoid arthritis, systemic lupus erythematosus, Sjogren’s syndrome, multiple sclerosis and inflammatory bowel disease, where it is associated with insertion/deletion mutations creating an Sp1 binding site that upregulates IRF5 expression.[19] IRF5 is therefore an attractive target for treatment of these disorders and could certainly be relevant for AS. The potential benefits of modulating IRF5 expression, its post-translational modification and/or functional interactions with its protein partners for therapeutic benefit have been extensively discussed elsewhere.[17] Increased binding of the chromatin reader BRD4, also to the AS-protective allele G (Bromodomain Containing 4, see Supplementary Table 1), reinforces the likely contribution to AS of factors involved in transcriptional control and epigenetic regulation at *RUNX3.[*20]

The approach we have adopted here is a first step towards understanding the mechanisms underlying one of the strongest genetic associations with AS and the role of *RUNX3* in its pathogenesis. In this paper and in previous work [7] we have demonstrated the impact of AS-associated SNPs on TF binding to the enhancer-like region upstream of *RUNX3* and on RUNX3 expression in CD8+ T-cells.[7] Further clarification of the pathways involved is needed but these results suggest that *RUNX3* related pathways and IRF5 represent important potential therapeutic targets for the treatment of AS.

## KEY MESSAGES

### What is already known about this subject?

- Ankylosing spondylitis (AS) has a strong genetic contribution.
- *RUNX3*, a transcription factor (TF) involved in CD8+ T-cell development and differentiation, has one of the strongest genetic associations with AS
- *rs4648889*, a single nucleotide polymorphism located upstream of the promoter influences *RUNX3* expression in CD8+ T-cells by altering transcription factor binding

### What does this study add?

- We identified many proteins, including transcription factors bound to the region encompassing *rs4648889* using a DNA pull-down approach coupled with quantitative Mass Spectrometry
- IRF5, IKZF3 and members of the gene repressor Nucleosome Remodelling Deacetylase (NuRD) complex exhibit differential binding between the AS-risk and AS-protective alleles.

### How might this impact on clinical practice or future developments?

- Differential binding of transcription factors at *rs4648889* suggests that regulation of *RUNX3* expression by IRF5 and the NuRD complex are important in the pathogenesis of AS
- RUNX3-related pathways and IRF5 represent important potential therapeutic targets for AS.

## AUTHOR CONTRIBUTIONS

MV, AC, RF, BPW and CC conceived and designed the experiments. MV, AC and RF performed the experiments. MV and RF analysed the data. MV and PW drafted the manuscript and all the authors revised the final version prior submission.

## ACKNOWLEDGMENTS

MS data was generated in the Discovery Proteomics Facility headed by Roman Fischer, as part of the TDI biological mass spectrometry laboratory under Benedikt M. Kessler.

We thank Dr. Adam Cribbs for his constructive comments to the manuscript.

---

**Supplementary Table1.**
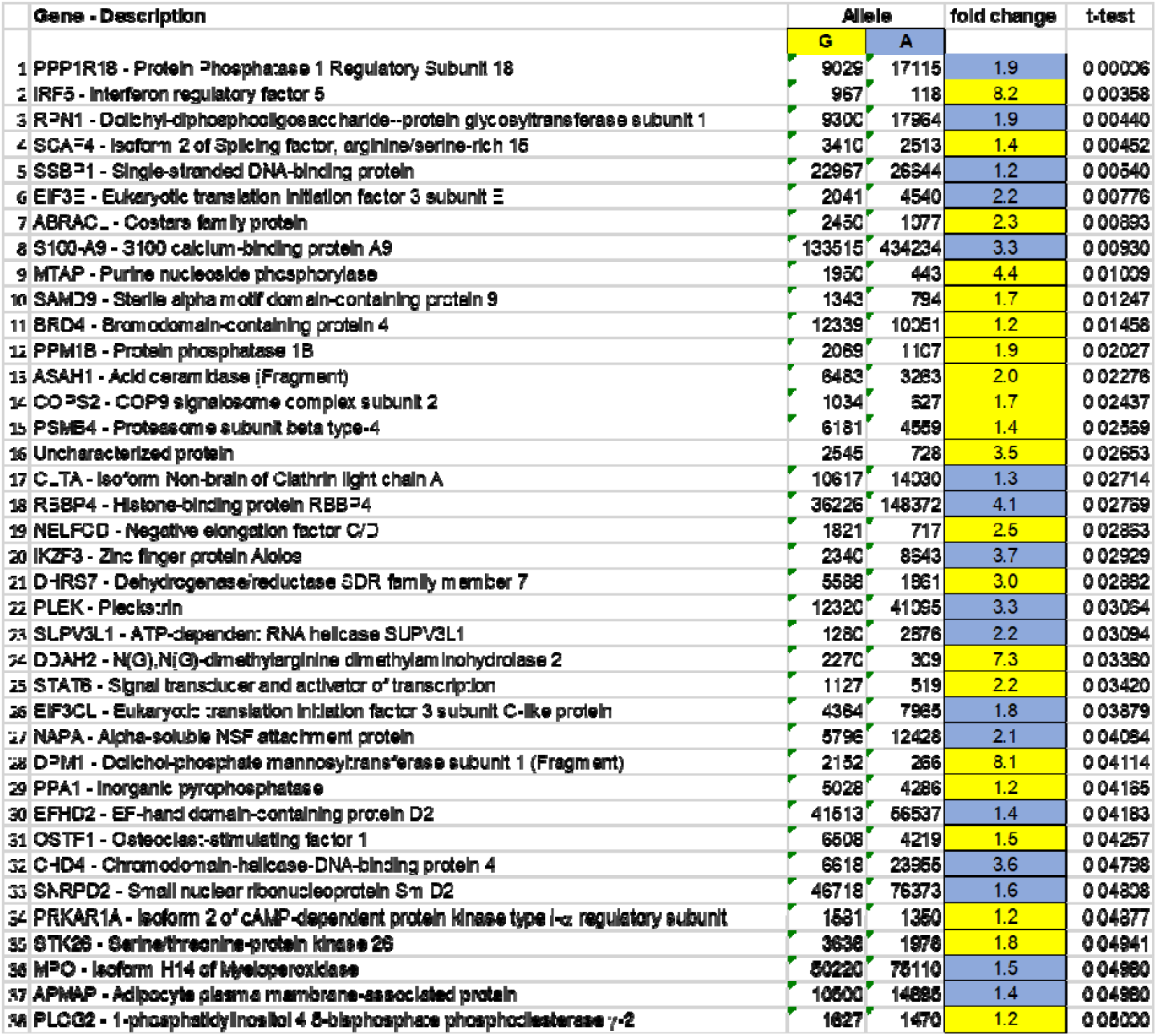
List of proteins with significantly differential binding affinity (p<0.05) to probes corresponding to the two alleles of *rs4648889*. Label-free quantification results for G (protective) and A (AS-risk) alleles, expressed as average, obtained from three independent experiments. Fold change and t-test are also shown. Proteins showing greater abundance with the “G” allele are highlighted in yellow while in light blue those for the “A” allele.

**Supplementary Table 2.**
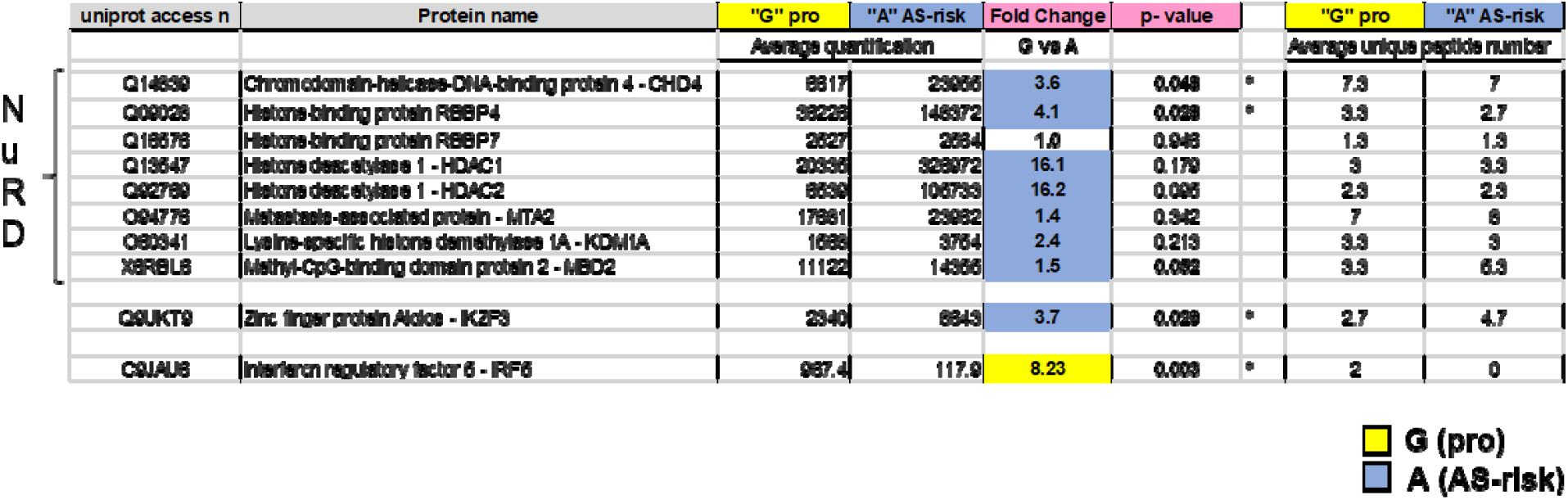
Label free quantification of IRF5, IKZF3 and NuRD complex factors identified with M/S approach. Label free quantification after median subtraction of IKZF3, IRF5 and the NuRD complex factors obtained from three independent experiments. NuRD proteins showed greater abundance with the “A” allele, highlighted in light blue. IRF5 showed significantly more abundance with the “G” allele. The unique peptide number is also shown (as average of three experiments). *Significant p-value.

**Supplementary Table 3.**
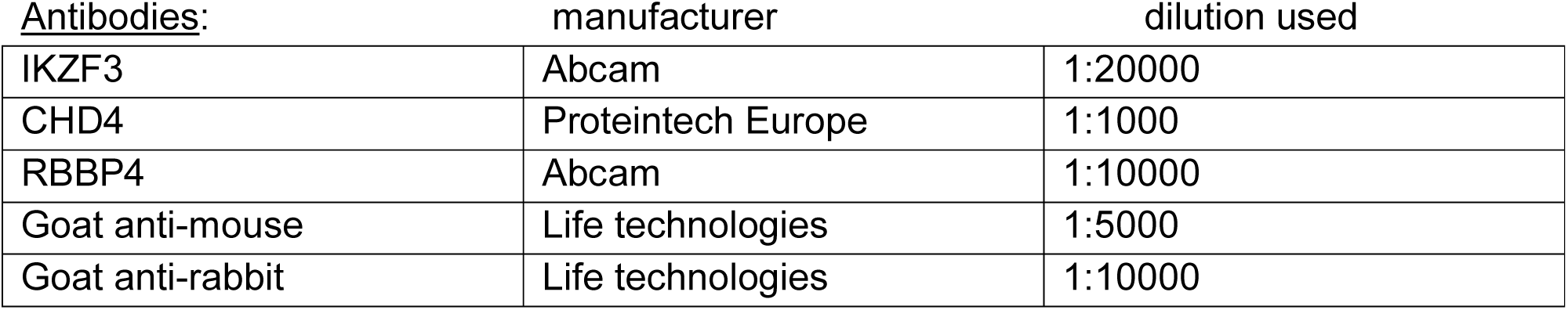
List of antibodies used for Western blot analysis.

**Supplementary Table 4.**
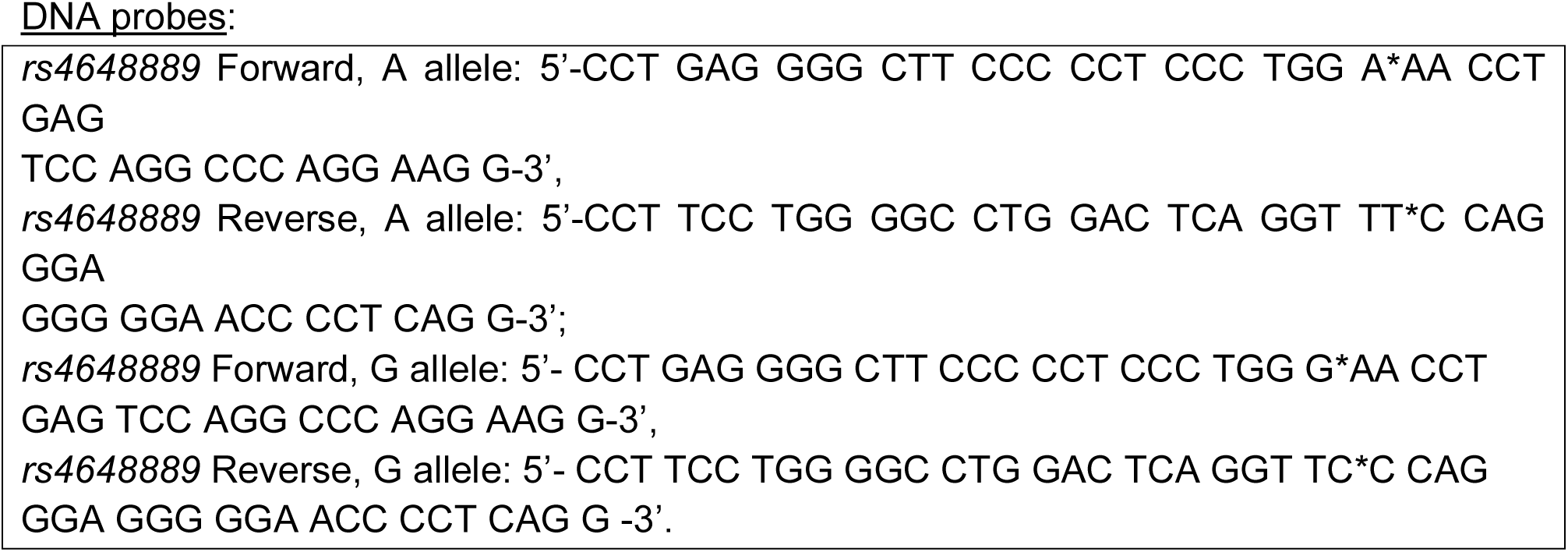
List of DNA probes for EMSA and pull-down experiments. Asterisks locate SNP *rs4648889*.

## METHODS

### DNA-affinity capture

100nM anti-sense single-stranded oligonucleotides (50bp) were 3’-end biotinylated, mixed and annealed at room temperature (RT) for 1hr with the sense oligonucleotide. Streptavidin-coated beads (Dynabeads M-280, catalogue number 11205D, Thermo Fisher Scientific, Waltham, Massachusetts, USA) were equilibrated with six washes with wash buffer (10.0 mM Tris-HCl pH 7.4, 2.0 M NaCl, 1mM EDTA). Biotin-labelled DNA was incubated with streptavidin-coated beads for 1 hour at RT, on a rotary wheel. Three successive washes were performed to eliminate the unbound biotinylated-DNA. Nuclear extract (NE) (500μg) from CD8+T-cells was pre-incubated on ice for 20 min in mobility shift essay (EMSA) binding buffer (100mM Tris, 500mM KCl, 10mM DTT; pH 7.5) and then incubated with beads for 1 hour at 21°C on a rotary wheel. Beads were then stringently washed six times: once with 500ul of EMSA binding buffer, three times with wash buffer + 0.1% Tween20, twice with 50mM NH_4_HCO_3_. Beads were then re-suspended in sample buffer containing 10mM EDTA, 1% SDS and Benzonase (1U) for 60 minutes, at RT on a rotary wheel, in order to separate the DNA-protein complexes from the beads. Magnetic separation was used to separate the biotinylated nucleic acid from the supernatant. Benzonase was used to eliminate DNA and other contaminants to ensure that only the protein complexes were collected. Overview of the experimental approach is showed in Supplementary Figure 1.

### Liquid chromatography mass spectrometry

Protein samples were prepared for mass spectrometric analysis by tryptic digest as described previously [1]. Briefly, proteins were reduced and alkylated (DTT/Iodoacetamide) before digest with Trypsin (Promega) and desalting of peptides using C18 material (Sola, Thermo-Fisher). Peptides were then analysed on a nano LC-MS/MS platform consisting of Q-Exactive mass spectrometer and nano UPLC (both Thermo Fischer) [2]. Chromatographic separation of peptides was achieved on a easyspray column (75umx 500mm) using a gradient from 5% DMSO in 0.1% formic acid/5% Acetonitrile to 5%DMSO in 0.1% formic acid in 35% Acetonotrile. MS parameter were used as described earlier. Quantitative data are derived from the number of MS/MS spectra per peptide (spectral counting) or the integrated peak area of the ion chromatogram of a specific peptide as reported by ProgenesisQI (Waters, version 2.0) using default parameters. Protein IDs were generated with Mascot at 1% FDR and peptide score cut-off at 20 against a Uniprot human database. All proteomics data is publicly available through the Pride consortium [3].

### Data visualization, STRING and Gene Ontology (GO) analysis

Proteomics data were visualized with Qlucore Omics Explorer (version 3.7) to obtain principal component analysis (PCA) and unsupervised hierarchical clustering. Cytoscape (v3.7.0) was used to analyse the MS data in defining Gene Ontology (GO) categories and sub-groups (http://geneontology.org/). The Reactome [4] database was used to integrate pathway analysis. The R package was used to create Volcano plots. String (v 11.0), a database of known and predicted protein-protein interactions (PPI), was used to define the PPI enrichment of the differentially abundant proteins identified.

### Epigenetic database interrogation

A combination of epigenetic data from the ENCODE [5] and Roadmap Epigenomics Projects [6] was used to analyse the region encompassing SNP *rs4648889*, in particular ChIP-seq peaks for IKZF and CHD4 transcription factor (TF) on lymphoblastoid cell lines.

### CD8+ T-cells□isolation and nuclear extract preparation

CD8+ T-cells were isolated from peripheral blood mononuclear cells (PBMCs) from buffy coat using a CD8+ T-cell isolation kit (Miltenyi, Bisley, Surrey, UK). CD8+ T-cells were re-suspended at 1×10^6^/mL in pre-warmed Roswell Park Memorial Institute (RPMI) medium supplemented with 10% foetal bovine serum, penicillin/streptomycin and L-glutamine. After 4 hours resting the cells were then harvested.

### Electrophoretic mobility gel shift assay (EMSA)

EMSA were performed as previously described [12]. Briefly, the DNA probes were mixed and annealed at room temperature for 1□hour. For super shift assays, 5μg of nuclear extract (NE) obtained from Jurkat cells were first incubated (20 min) with a specific antibody, then the NE-antibody complex was incubated (20 min) with biotinylated DNA and run on retardation gels. Full list of antibodies and DNA probes are available in Supplementary Table 3 and 4.

### Western blot assay

Eluted samples were separated by SDS–PAGE, transferred onto nitrocellulose membranes (Bio-Rad Laboratories Berkeley, California, USA), and incubated overnight at 4°C variously with primary antibodies against IKZF3, CHD4, RBBP4. Appropriate horse-radish peroxidase-conjugated secondary antibody was used and signals detected with enhanced chemiluminescence (ECL) method (Thermo Fisher Scientific, Waltham, Massachusetts, USA).

### ChIP-qPCR

ChIP was performed using ABI ViiA 7 PCR device (Applied Biosystems, Paisley, UK) using the iDeal ChIP-seq kit for Transcription Factors (catalogue number C01010055, Diagenode, Liege, Belgium). For each ChIP, 2.5×10^6^ CD8+ T-cells were used. Three independent qPCR experiments were performed using allele-specific primers for *rs4648889* (primers sequence available here [12]). We used CD8+ T-cells (n=3) of known genotype (heterozygous for *rs4648889*) from buffy coat blood cones to compare the impact of the AS-risk and -protective alleles on relative enrichment. We normalized all of our ChIP-qPCR data against a 1% input control as per manufacturer instructions. Data were visualized with Prism version 8.0.2. The following ChIP grade antibodies were used: RBBP4 (catalogue number ab79416, Abcam, Cambridge, UK), CHD4 (catalogue number 14173-1-AP, Proteintech Europe, Manchester, UK), IKZF3 (catalogue number ab139408, Abcam, Cambridge UK), IRF5 (catalogue number E1N9G Rabbit mAb #13496) and IgG (catalogue number K02041008, Diagenode, Liege, Belgium).

### Patient and Public Involvement

The database used in the study was developed with direct involvement of the National Ankylosing Spondylitis Society through its members, trustees and medical advisory committee. All aspects of this work are regularly reported to the members of the Society through its bimonthly newsletter and annually at the AGM and members’ day.

